# Trisomy 21 increases microtubules and disrupts centriolar satellite localization

**DOI:** 10.1101/2021.10.25.465808

**Authors:** Bailey L. McCurdy, Cayla E. Jewett, Alexander J. Stemm-Wolf, Huy Nguyen Duc, Molishree Joshi, Joaquin M. Espinosa, Rytis Prekeris, Chad G. Pearson

## Abstract

Trisomy 21, the cause of Down syndrome, causes a 0.5-fold protein increase of the chromosome 21-resident gene Pericentrin (PCNT) and reduces primary cilia formation and signaling. Here we investigate the mechanisms by which PCNT imbalances disrupt cilia. Using isogenic RPE-1 cells with increased chromosome 21 dosage, we find PCNT protein accumulates around the centrosome as a pericentrosomal cluster of enlarged cytoplasmic puncta that localize along and at MT ends. Cytoplasmic PCNT puncta impact the intracellular MT trafficking network required for primary cilia, as the PCNT puncta sequester cargo peripheral to centrosomes in what we call pericentrosomal crowding. The centriolar satellite proteins, PCM1, CEP131 and CEP290, important for ciliogenesis, accumulate at sites of enlarged PCNT puncta in trisomy 21 cells. Reducing PCNT when chromosome 21 ploidy is elevated is sufficient to decrease PCNT puncta, reestablish a normal density of MTs around the centrosome, restore ciliogenesis to wild type levels and decrease pericentrosomal crowding. A transient reduction in MTs also decreases pericentrosomal crowding and partially rescues ciliogenesis in trisomy 21 cells, indicating that increased PCNT leads to defects in the microtubule network deleterious to normal centriolar satellite distribution. We propose that chromosome 21 aneuploidy disrupts MT-dependent intracellular trafficking required for primary cilia.

**TOC:** McCurdy *et al* explore why elevated Pericentrin in trisomy 21 negatively impacts primary cilia formation and signaling. They find that elevated Pericentrin produces more pericentrosomal puncta that associates with and increases microtubules. Changes to Pericentrin and microtubules mislocalizes centriolar satellites in a pericentrosomal crowd.

## Introduction

Primary cilia are signaling platforms required for development and homeostasis, and ciliopathies are genetic disorders affecting genes that impact ciliary function (Breslow and Holland, 2019; Nigg and Raff, 2009; Reiter and Leroux, 2017). Down syndrome (DS), or trisomy 21, is a distinct genetic condition caused by an extra copy of human somatic autosome 21 (HSA21) that also negatively impacts development (Hattori et al., 2000). Consistent with the phenotypic overlap between pathologies associated with DS and ciliopathies, trisomy 21 produces primary cilia and cilia associated signaling defects (Currier et al., 2012; Galati et al., 2018; Ripoll et al., 2012; Roper et al., 2006).

Primary cilia assemble from the centrosome comprised of two centrioles surrounded by a dense protein matrix called the pericentriolar material (PCM) (Bettencourt-Dias et al., 2011; Bornens and Azimzadeh, 2007; Gould and Borisy, 1977; Nigg and Stearns, 2011; Sorokin, 1962). The mother centriole nucleates and organizes the primary cilium (Breslow and Holland, 2019). The centrosome PCM organizes cytoplasmic microtubules (MTs) required for trafficking cargoes to and from centrosomes, including cargoes required for cilia (Kim et al., 2004; Kubo et al., 1999; Sorokin, 1962; Sung and Leroux, 2013; Woodruff et al., 2014). In addition to the centrosome, the golgi apparatus nucleates MTs responsible for cell migration in interphase cells (Efimov et al., 2007; Gavilan et al., 2018). Vesicles moving from the golgi apparatus to the centrosome are critical for early events in cilia formation (Follit et al., 2006; Shakya and Westlake, 2021). In the case of primary cilia formation during interphase, MT minus ends emanate from the centrosome PCM, establishing a distribution hub for MT-motor driven cargoes to and from the cilium (Dammermann and Merdes, 2002; Kubo *et al*., 1999). To ensure accurate cargo translocations, the number and stability of MTs are tightly controlled (Doxsey, 2001; Vertii et al., 2016; Woodruff *et al*., 2014).

Within the trafficking milieu required for cilia formation and function are centriolar satellites. Centriolar satellites are 70-100 nm non-membrane bound granule structures that localize around the centrosome and move along MTs (Bernhard and de Harven, 1960; Dammermann and Merdes, 2002; Hori and Toda, 2017; Kubo *et al*., 1999; Odabasi et al., 2019). Centriolar satellites positively and negatively modulate primary cilia through protein transport, degradation, and sequestration (Odabasi *et al*., 2019; Prosser and Pelletier, 2020; Tollenaere et al., 2015). The Pericentriolar Material-1 (PCM1) centriolar satellite scaffold protein ensures centriolar satellite integrity and is required for ciliogenesis (Gupta et al., 2015; Wang et al., 2016). PCM1 loss disassembles centriolar satellites, reduces PCM, and disrupts primary cilia (Dammermann and Merdes, 2002; Keryer et al., 2011; Odabasi *et al*., 2019; Prosser and Pelletier, 2020; Quarantotti et al., 2019). The dynamic interplay between centrosomal PCM and centriolar satellites is instrumental for primary cilia, yet how it is controlled and how it is disrupted in disease remains poorly understood.

Pericentrin (PCNT) is a centrosome PCM scaffold protein essential for ciliary protein recruitment and MT nucleation (Delaval and Doxsey, 2010; Sung and Leroux, 2013; Woodruff *et al*., 2014). Along with CDK5RAP2/CEP215, PCNT recruits *γ*-tubulin to the centrosome (Dictenberg et al., 1998; Fong et al., 2008). PCNT also interacts with PCM1 and supports MT motor-dependent trafficking to and from centrosomes in particles that variably associate with centriolar satellites (Dammermann and Merdes, 2002; Doxsey et al., 1994; Galati *et al*., 2018; Young et al., 2000). PCNT is not necessary for interphase centrosome MT organization but is essential for mitotic spindle assembly (Dammermann and Merdes, 2002; Gavilan *et al*., 2018; Zimmerman et al., 2004). However, PCNT forms a complex with intraflagellar transport proteins, and its loss or overexpression disrupts IFT20 localization to the centrosome and primary cilia formation (Galati *et al*., 2018; Jurczyk et al., 2004). The precise roles for PCNT in promoting cilia formation are unknown. Moreover, PCM1 and PCNT interactions promote centrosome PCM assembly (Dammermann and Merdes, 2002; Li et al., 2001). The PCNT gene is on HSA21 and protein levels are elevated in DS (Galati *et al*., 2018; Salemi et al., 2013). Using heterologous and mosaic DS cell lines, elevated PCNT was found to disrupt primary cilia (Galati *et al*., 2018). Elevated PCNT in trisomy 21 produces enlarged centrosomes and cytoplasmic PCNT puncta that behave as phase dense particles (Galati *et al*., 2018; Jiang et al., 2021). As puncta enlarge, the movement of PCNT along MTs is suppressed, as is the localization of factors required for ciliogenesis (IFT20) and ciliary dependent signaling (Smoothened) (Galati *et al*., 2018). A mechanistic understanding for how elevated and enlarged PCNT puncta control MTs, centriolar satellite localization, and primary cilia formation is still required.

## Results and Discussion

### Disrupted primary cilia in increased HSA21 isogenic lines is PCNT dependent

To study the controlled effects of HSA21 dosage on primary cilia, we utilized isogenic RPE1 cell lines (disomy 21; D21) engineered with either one (trisomy 21; T21) or two (tetrasomy 21; Q21) extra copies of HSA21 (Lane et al., 2014). Primary cilia frequency was quantified in cells incubated for 24 hours in reduced serum medium where T21 or Q21 cells were co-cultured with D21 cells to control for confluency and non-cell autonomous effects (Fig. 1, A and B; and Fig. S1 A). Cilia frequency was reduced in an HSA21 dose dependent manner (32% and 56% reduction for T21 and Q21; Fig. 1 B). Primary cilia length was not changed and Q21 cells were slightly larger than D21 and T21 cells (Fig. S1, C and D). Total cellular protein levels of the HSA21 resident gene, PCNT, in T21 and Q21 cells were increased to 1.3- and 2-fold, respectively (Fig. S1, B and G). Importantly, PCNT levels surrounding the centrosome in unciliated T21 and Q21 cells were 42% and 31% greater than their ciliated cell counterparts, respectively (Fig. 1 D). Thus, elevated PCNT is associated with decreased cilia formation. However, there is no strict PCNT threshold across the three cell lines above which cilia do not form. This suggests that cells compensate for elevated PCNT and/or that increased HSA21 dosage suppresses some of the negative impacts elevated PCNT has on cilia. To confirm that it is elevated PCNT that impacts cilia frequency, PCNT was reduced in T21 and Q21 cells. Reduction of PCNT in T21 and Q21 cells to approximately D21 levels using siRNA increased cilia frequencies to 92% and 50% of D21 levels, respectively (Fig. 1 E). Genetic ablation of a single PCNT allele in T21 cells using CRISPR-Cas9 (leaving cells with 2n PCNT) also increased the mean cilia frequency to 121% of D21 levels (Fig. 1 E; and Fig. S1, E-I). One possibility for the incomplete recovery in the siRNA experiments is because cells were coincidentally depleted for PCNT and induced for ciliogenesis such that ciliating cells do not have the complete 24 hours with normal PCNT levels. Consistent with this, the genetic ablation studies completely rescued the cilia frequency. In summary, a dose-dependent increase in HSA21 and PCNT inhibits primary cilia formation in isogenic RPE1 cells.

**Figure 1:**
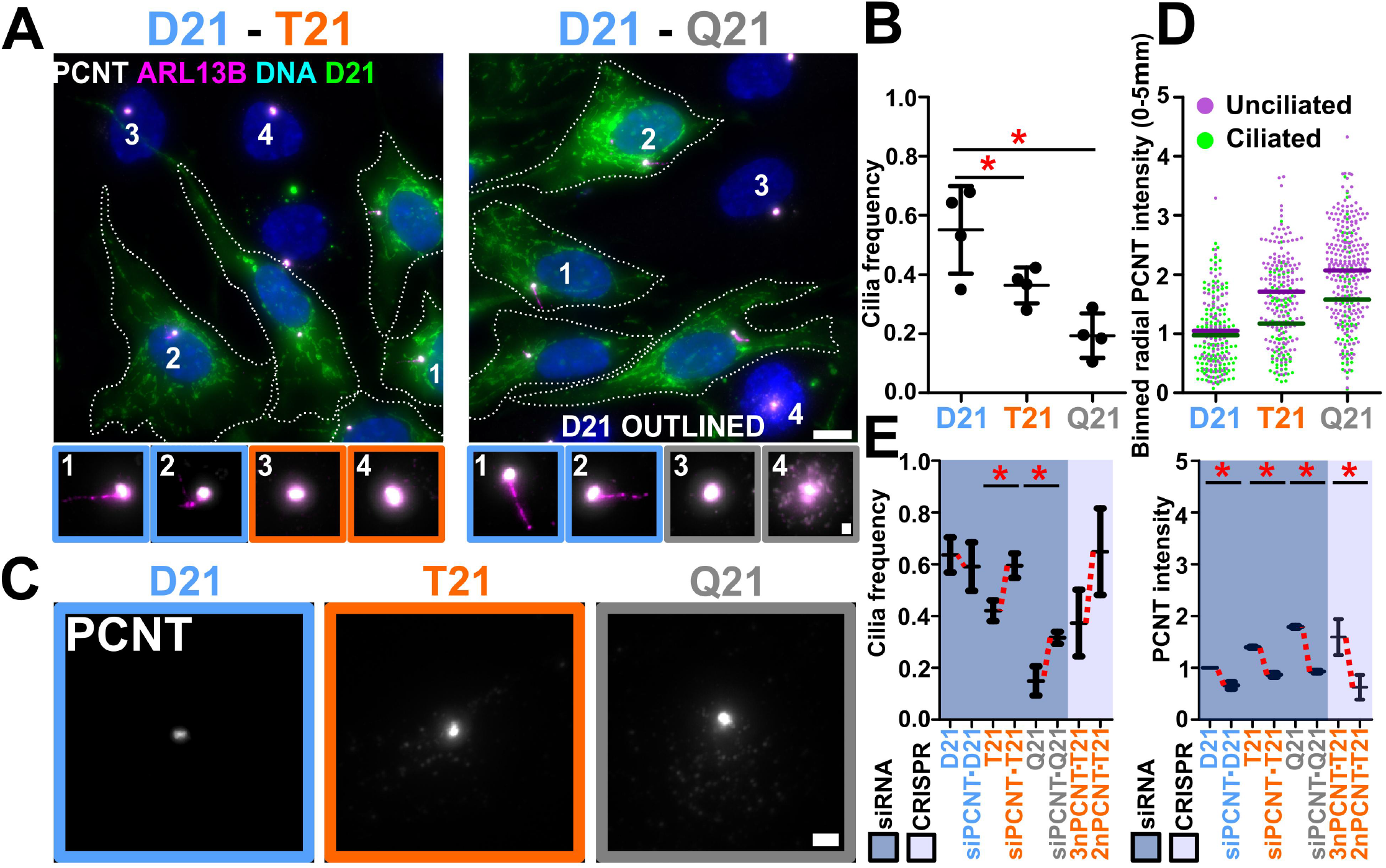
Elevated chromosome 21 dosage and PCNT disrupt primary cilia formation. **(A)** Left panel, co-cultured disomy 21 (D21) and trisomy 21 (T21) RPE1 cells stained for centrosomes (Pericentrin (PCNT); grayscale), cilia (ARL13B; magenta) and DNA (Hoescht33258; blue). Right panel, analogously stained co-cultured D21 and Q21 cells. D21 cells were CFSE stained (green) and outlined. Cell number labels are referenced in bottom panels. Scale bars, 10 μm and 1 μm for insets. **(B)** Cilia frequency decreases with HSA21 ploidy. Mean±SD. *p < 0.05 (Table 1). **(C)** PCNT florescence at the centrosome (PCNT; grayscale) in D21, T21, and Q21 RPE1 cells. Scale bar, 2 μm. **(D)** PCNT levels increase with HSA21 ploidy and unciliated cells. Intensity values normalized to D21 average. Data points represent PCNT fluorescence intensity within a 5 μm radius of the centrosome. **(E)** Reducing PCNT via siRNA or CRISPR-Cas9 knockout of one allele of PCNT increases cilia frequencies in T21 and Q21 RPE1 cells. Changes in mean values indicated with red lines. Intensity values normalized to D21 average. Data points represent PCNT fluorescence intensity within a 5 μm radius of the centrosome. Mean±SD. * p < 0.05 (Table 1).

### Elevated PCNT forms enlarged puncta encircling the pericentrosomal region that is associated with reduced cilia

To understand how elevated PCNT inhibits cilia formation, we investigated differences in PCNT protein distribution at and around the centrosome relative to total cellular levels. In addition to increased PCNT at the centrosome, enlarged PCNT puncta surround the centrosome (Fig. 2 A; (Galati *et al*., 2018)). The radial distribution of total PCNT fluorescence intensity around the centrosome was quantified and normalized to ciliated D21 PCNT fluorescence intensity levels (Fig. 2, B and C; and Fig. S2 A). The peripheral non-centrosomal PCNT surrounding the centrosome (1.2-5.0 μm) represents 45-53% of total cellular PCNT intensity (Fig. S2 B). Although T21 and Q21 PCNT levels were elevated in all regions, PCNT is increased by 1.6- and 1.9-fold at the centrosome and 1.5- and 2.3-fold in regions surrounding the centrosome in unciliated T21 and Q21 cells as compared to their ciliated cell counterparts (Fig. 2 C; and Fig. S2 A). The population of PCNT around the centrosome is comprised of both large and small puncta (Fig. 2, A and D). Enlarged puncta are greater than 0.05 μm^2^ and commonly associated with MTs (see below) (Fig S2 I). Unciliated cells had more enlarged PCNT puncta than ciliated cells (Fig. 2 D), consistent with the hypothesis that their accumulation inhibits primary cilia formation. Moreover, decreasing PCNT levels with siRNA or CRISPR-Cas9 reduces PCNT fluorescence intensity both at and around centrosomes (Fig. 2, E and F). This suggests increases to PCNT surrounding centrosomes causes fewer cilia.

**Figure 2:**
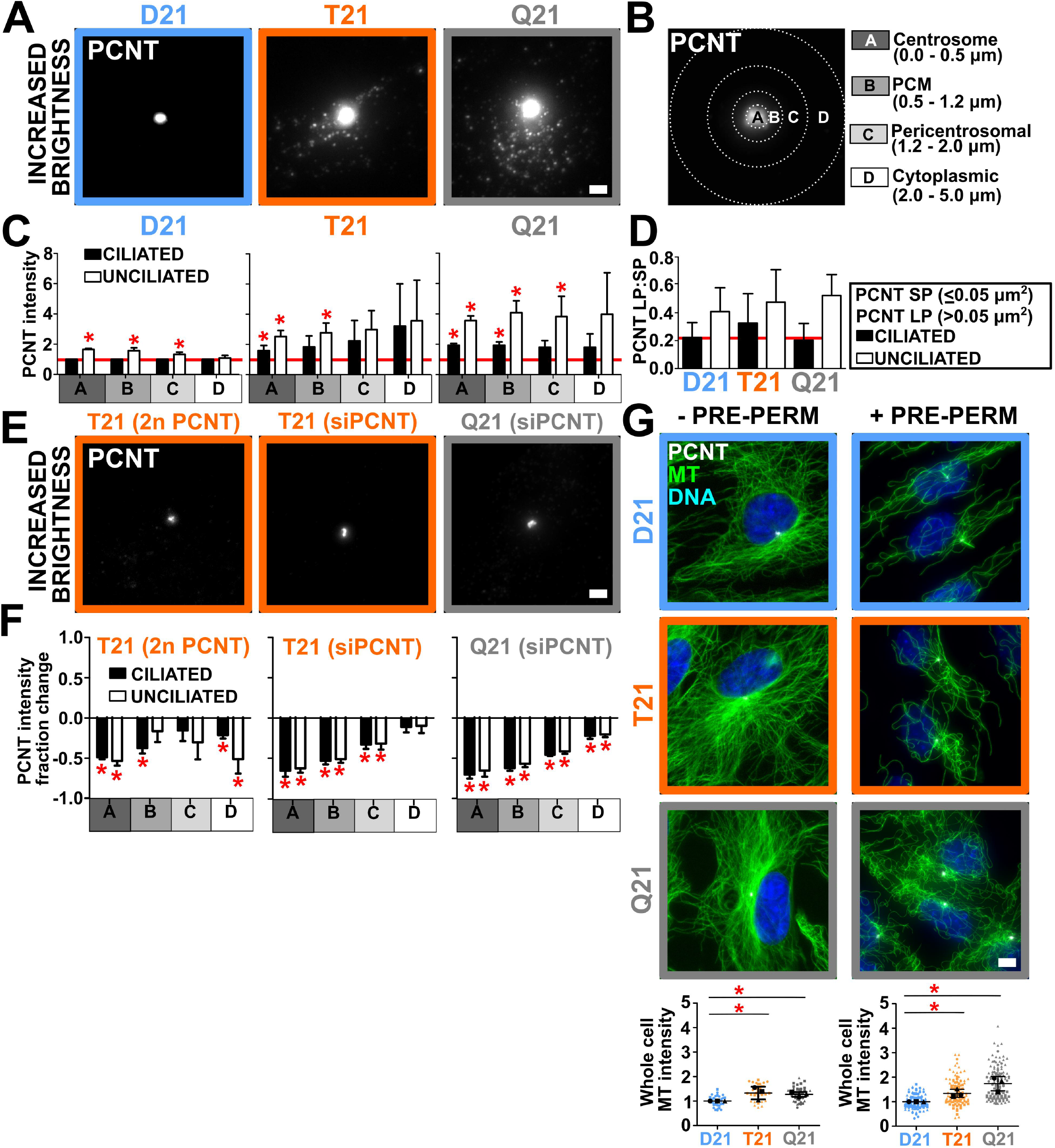
Elevated PCNT forms large puncta peripheral to the centrosome that disrupt cilia formation. **(A)** PCNT fluorescence (PCNT; grayscale) in D21, T21, and Q21 RPE1 cells. Brightness increased by a factor of 2.5. Scale bar, 2 μm. **(B)** Centrosome (A), pericentriolar (B), pericentrosomal (C), and cytoplasmic (D) regions for binned radial fluorescence intensity analysis from the centroid of the centrosome. **(C)** D21, T21, and Q21 cell regional binned PCNT fluorescence intensities. Intensity values normalized to ciliated D21 average (indicated with red line). Statistical comparisons made to ciliated D21 averages. Mean±SD. *p < 0.05 (Table 1). **(D)** PCNT large puncta (LP):small puncta (SP) ratios in ciliated and unciliated D21, T21, and Q21 cells. Mean±SD. **(E)** PCNT fluorescence (PCNT; grayscale) in PCNT CRISPR-Cas9 knockout line (T21 (2n PCNT)) and cells treated with PCNT siRNA (T21 (siPCNT), Q21 (siPCNT)). Scale bar, 2 μm. **(F)** Reducing PCNT via CRISPR-Cas9 knockout of one allele in T21 and siRNA in T21 and Q21 RPE1 cells reduces PCNT intensity at and peripheral to the centrosome. Statistical comparisons made to relative control intensities. Mean±SD. *p < 0.05 (Table 1). **(G)** Left panels, D21, T21 and Q21 RPE1 cells stained for MTs (DM1A; green), PCNT (PCNT; grayscale), and DNA (Hoescht33258; blue). MTs in these images were fixed without pre-permeabilization step. Scale bar, 5.5 μm. Right panels, D21, T21 and Q21 RPE1 cells stained for MTs (DM1A; green), PCNT (PCNT; grayscale), and DNA (Hoescht33258; blue). MTs in these images were fixed with pre-permeabilization step. Scale bar, 5.5 μm. Left graph, D21, T21, and Q21 whole cell non-pre-permeabilized MT fluorescence intensities. Right graph, D21, T21, and Q21 whole cell pre-permeabilized MT fluorescence intensities. Intensity values normalized to D21 average. Statistical comparisons made to D21 averages. Mean±SD. *p < 0.05 (Table 1).

### Elevated PCNT puncta increase cytoplasmic MTs

Cytoplasmic PCNT puncta arise from either self-assembly or from the centrosome and localize in the pericentrosomal space. We suggest that these puncta are analogous to the cytoplasmic MT organizing centers found when centrioles, centrosomes and the golgi apparatus are disrupted (Gavilan *et al*., 2018). To test whether the elevated levels of centrosomal and pericentrosomal PCNT increase total cellular MTs, we measured the effects of increased HSA21 dosage on MTs. Total cell MT intensities were elevated with increased HSA21 dosage (Fig. 2G; and Fig. S2, C and D). Because MTs of interphase RPE1 cells nucleate from both the centrosome and golgi apparatus (Efimov *et al*., 2007; Gavilan *et al*., 2018), we next tested whether elevated MT intensity was associated with the golgi apparatus (Fig. S2 E). In contrast to the centrosome and pericentrosomal regions, only a minor increase in MT intensity was observed to overlap with the golgi apparatus marked with IFT20 (Fig. S2 E). The contribution of golgi apparatus MTs relative to whole cell MTs was not significantly elevated across D21, T21, and Q21 cells (Fig. S2 E). This suggests that even though trafficking from the golgi apparatus to the centrosome may be disrupted (Galati *et al*., 2018), MTs are predominantly elevated at the centrosome and pericentrosomal regions, and that elevated PCNT plays a role in increased MT nucleation and / or stability.

To test whether MTs are more stable in T21 and Q21 cells, cells were exposed to either a 5-minute cell pre-permeabilization or an acute 10 minute cold treatment (cold MT depolymerization (CD)), both of which disassemble dynamic MTs (Fig. 2 G; and Fig. S2, F and G). The 5-minute pre-permeabilization prior to fixation resulted in increased MT densities in T21 and Q21 cells when compared to D21 cells receiving the same treatment (Fig. 2 G). Moreover, consistent with T21 and Q21 having greater MT stability, increasing MT densities were found in T21 and Q21 cells as compared to D21 cells after a 10 minute CD (Fig. S2, F and G). Moreover, the percent decrease of MTs after a 10-minute CD was reduced in T21 and Q21 cells compared to D21 cells (Fig. S2 G). Together, these results suggest that MT stability is increased in T21 and Q21 cells.

To address whether MT post-translational modifications (PTMs) contribute to increased MT stability, tubulin and tubulin PTM densities were quantified (Fig. S2 H). Tubulin density was slightly increased in Q21 cells, however, this increase was not enough to account for the differences in the whole cell MT intensities that were observed in these cells (Fig S2 H). Moreover, detyrosinated tubulin was increased in Q21 cells and acetylation and glutamylation remained unchanged under the conditions tested (Fig. S2 H). In summary, increased MT stability in T21 and Q21 cells is likely independent of MT PTMs.

### Enlarged PCNT puncta increase MTs and inhibit ciliogenesis

We next investigated whether elevated PCNT and PCNT puncta in T21 and Q21 cells contribute to the increased MT density at and around the centrosome. Indeed, both regional and whole cell MT densities using pre-extraction fixation methods were increased in both unciliated cells and cells with elevated HSA21 dosage (Fig. 3, A and B; and Fig. S3, A and B). D21 and Q21 cells exhibited significant positive correlation coefficients when comparing whole cell PCNT and MT intensities (Fig. S3 B). Coincident with increased PCNT intensities (Fig. 2 C; and Fig. S2 A), MTs in T21 and Q21 cells were elevated in all regions surrounding centrosomes (Fig. 3 B; and Fig. S3 A). The levels of PCNT interacting partners, CEP215 and *γ*-tubulin, were also increased at and surrounding centrosomes (Fig. S3, C-E). In T21 and Q21 cells, an increase in *γ*-tubulin and PCNT colocalization is observed at free MT ends (Fig. S3 F). This suggests that the increase in PCNT associated with HSA21 dosage and unciliated cells increases MT-nucleation factors and MTs. Because a larger increase in MTs is observed in the pericentrosomal region in unciliated cells, we hypothesize this PCNT and MT population is responsible for disrupting cilia formation.

**Figure 3:**
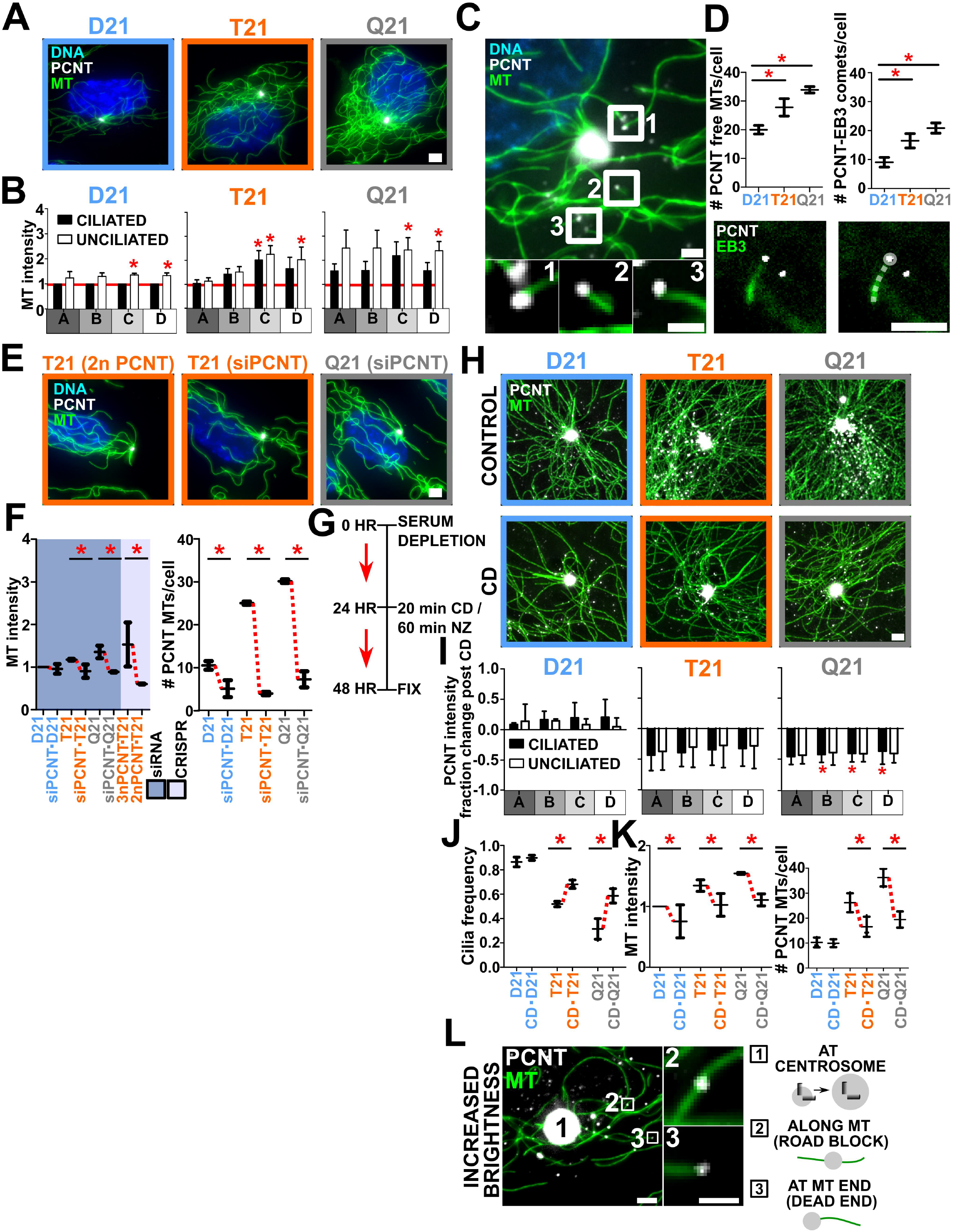
Large PCNT puncta nucleate free microtubules that inhibit ciliogenesis. **(A)** D21, T21 and Q21 RPE1 cells stained for MTs (DM1A; green), PCNT (PCNT; grayscale), and DNA (Hoescht33258; blue). A pre-permeabilization step was included prior to fixation. Scale bar, 3 μm. **(B)** D21, T21, and Q21 cell regional binned MT fluorescence intensities. Intensity values normalized to ciliated D21 average (indicated with red line). Statistical comparisons made to ciliated D21 averages. Mean±SD. *p < 0.05 (Table 1). **(C)** PCNT localizes to MT ends. Unciliated D21 RPE1 cell stained for MTs (DM1A; green) and PCNT (PCNT; grayscale). Labels are referenced in bottom panels. Scale bars, 2 μm and 0.75 μm for insets. **(D)** Top, Hsa21 dosage increases PCNT colocalization at MT ends and PCNT nucleated MTs. Bottom, RPE1 cell labeled with EB3-mNeon and PCNT (PCNT; grayscale). Right panel includes a line trace of the EB3 comet. Scale bars, 2.5 μm. Mean±SD. *p < 0.05 (Table 1). **(E)** Crispr-Cas9 knockout line (T21 (2n PCNT)) and cells treated with PCNT siRNA (T21 (siPCNT), Q21 (siPCNT)) stained for MTs (DM1A; green), PCNT (PCNT; grayscale), and DNA (Hoescht33258; blue). Scale bar, 3 μm. **(F)** Left, reducing PCNT via siRNA in T21 and Q21 RPE1or CRISPR-Cas9 knockout of one allele in T21 cells reduces MT intensities. Intensity values normalized to D21 average. Data points represent 5μm binned radial MT fluorescence intensities. Right, reducing PCNT via siRNA reduces PCNT nucleated MTs in D21, T21, and Q21 RPE1 cells. Mean±SD. *p < 0.05 (Table 1). **(G)** Timeline for cold-depolymerization (CD) and nocodazole (NZ) treatment experiments. **(H)** Top panels, D21, T21 and Q21 RPE1 cells stained for MTs (DM1A; green) and PCNT (PCNT; grayscale). Bottom panels, D21, T21 and Q21 RPE1 CD treated cells stained for MTs (DM1A; green) and PCNT (PCNT; grayscale). Scale bar, 1.5μm. **(I)** Cold-depolymerization reduces PCNT intensities at and peripheral to the centrosome in T21 and Q21 RPE1 cells. Statistical comparisons made to relative control intensities. Mean±SD. *p < 0.05 (Table 1). **(J)** Cold-depolymerization increases cilia frequencies in T21 and Q21 RPE1 cells. Mean±SD. *p < 0.05 (Table 1). **(K)** Left, cold-depolymerization reduces MT intensities in T21 and Q21 RPE1 cells. Intensity values normalized to D21 average. Data points represent 5μm binned radial MT fluorescence intensities. Right, cold-depolymerization reduces PCNT nucleated MTs in D21, T21, and Q21 RPE1 cells. Mean±SD. *p < 0.05 (Table 1). **(L)** PCNT puncta localize along MTs (2), and at MT ends (3). RPE1 cell labeled for microtubules (MTs) (DM1A; green) and PCNT (PCNT; grayscale). Scale bars, 2.5 μm and 0.75 μm for insets.

To test whether PCNT puncta are associated with free MT ends, the number of centrosome-free MTs associated with PCNT puncta was quantified (Fig. 3, C and D). An increase of 39% and 70% of centrosome-free MTs associated with PCNT puncta was observed in T21 and Q21 compared to D21 cells, respectively (Fig. 3 D). To determine whether these are sites of active MT growth, the number of growing MT plus ends (EB3 comets) associated with the enlarged PCNT puncta was quantified in D21, T21, and Q21 cells (Fig. 3 D). PCNT-associated EB3 comets increased with elevated HSA21 dosage (Fig. 3 D). This suggests that enlarged PCNT puncta in the pericentrosomal region act to organize MTs peripheral to the centrosome. Moreover, depletion of PCNT using siRNA or CRISPR-Cas9 reduced T21 and Q21 MT intensities and PCNT associated free-MTs to or below D21 levels (Fig. 3 F). Thus, enlarged cytoplasmic PCNT puncta promote increased MT growth in both unciliated cells and cells with elevated HSA21 dosage. We suggest that these MT populations repress cilia formation.

To investigate whether increased MTs caused by elevated PCNT contribute to primary cilia defects in T21 and Q21 cells, we reduced MTs using CD (Fig. 3, G-K). Following a 20-minute CD treatment and 24-hour recovery, MTs were reduced when compared to cells that did not receive the CD treatment (Fig. 3, H and K). Importantly, whole cell PCNT and PCNT at and surrounding T21 and Q21 centrosomes was reduced (Fig. 3, H and I; and Fig. S3 L). The relative mean primary cilia frequency increased by 31% and 86% in T21 and Q21 cells after CD treatment, respectively (Fig. 3 J). To address the model that PCNT-nucleated free MTs contribute to primary cilia defects in T21 and Q21 cells, the number of PCNT-associated free MTs post-CD was quantified (Fig. 3 K). CD reduced the number of PCNT associated free MTs in T21 and Q21 cells (Fig. 3 K). Thus, PCNT associated free MTs contribute to primary cilia defects.

### Enlarged PCNT puncta form along and at the ends of MTs

Because MT-dependent intracellular trafficking to and from the centrosome is disrupted by T21 (Galati *et al*., 2018), we proposed two non-mutually exclusive models by which enlarged PCNT puncta that associate with MTs inhibit intracellular trafficking (Fig. 3 L). First, PCNT puncta localized along MTs act as trafficking roadblocks. In this model, enlarged PCNT puncta either sterically block the movement of other cargoes or bind and sequester cargos as they encounter PCNT puncta. Cargo can translocate around objects on MTs (Can et al., 2014; Conway et al., 2012). Our prior work suggests that the trafficking of enlarged PCNT puncta is reduced along MTs (Galati *et al*., 2018), and PCNT-associated cargos no longer traffic efficiently. Second, given its MT nucleation capacity with CEP215 and *γ*-tubulin (Doxsey *et al*., 1994; Gavilan *et al*., 2018), PCNT puncta promote promiscuous free-MT nucleation at cytoplasmic puncta peripheral to the centrosome that generate trafficking dead-ends.

To visualize single MTs and PCNT puncta, cells were prepared using a 5-minute pre-permeabilization to focus on stable MTs and to facilitate the discernment of PCNT puncta associated with MTs (Fig. 2 G). Consistent with both models, PCNT puncta localize along MTs and at MT ends (Fig. 3 L). The average size of PCNT puncta along MTs increased in T21 and Q21 cells, while the average size of PCNT puncta at MT ends was unchanged in T21 and Q21 cells (Fig. S2 I). On average, 78% were associated with MTs (Fig. S2 J). Decreasing PCNT levels with siRNA or CRISPR-Cas9 reduces the number and size of enlarged PCNT puncta around the centrosome (Fig. 2, E and F). Because PCNT reduction in T21 and Q21 also increases ciliogenesis, we suggest that elevated and enlarged MT-associated PCNT puncta disrupt primary cilia formation.

To address the contribution of the elevated and enlarged PCNT puncta to the primary cilia defect, we treated cells with nocodazole (NZ) which disassembles MTs without disrupting large PCNT puncta (Fig. 3 G; and Fig. S3, G, I and J). Under these conditions, enlarged PCNT puncta remain while MTs are transiently depolymerized allowing us to test whether puncta are detrimental to cilia. Both NZ treatment and CD dissembled 60-85% of MTs in D21, T21, and Q21 cells (Fig. S3 J and K). MTs partially recover by 10 minutes and do not completely recover to their pre-depolymerization levels even 24 hours after CD or NZ treatment (Fig. S3 K). After transient NZ treatment and a 24 hour incubation in low serum conditions, the relative mean cilia frequency increased by 15% and 144% in T21 and Q21 cells, respectively (Fig. S3 F). Though mean increases in cilia post NZ are comparable to CD, large variability in cilia frequency post NZ but not CD suggests NZ rescues cilia less robustly than CD (% coefficient of variation (%CV) for NZ and CD treatment are: 10% and 2% for D21, 13% and 4% for T21, and 18% and 8% for Q21; Fig. 3 J; and Fig. S3 F). We attribute this to the negative effects of PCNT puncta on cilia formation that remain after NZ treatment but are dispersed after CD. Like PCNT reduction and CD treatment, MT intensities and PCNT nucleated free MTs were reduced in T21 and Q21 cells post-NZ treatment (Fig. S3 G). Following either PCNT siRNA, CD, or NZ treatment, the remaining PCNT-associated, free MTs were closer to the centrosome (Fig. S3 H). This suggests that a dynamic reorganization of PCNT puncta and MTs reestablishes the MT network required for primary ciliogenesis. However, unlike PCNT reduction or CD, NZ did not significantly reduce the elevated pericentrosomal PCNT in T21 and Q21 cells (Fig. S3 K). NZ treatment alters cellular MTs without disturbing enlarged PCNT puncta. PCNT nucleated centrosome-free MTs lead to MT-dependent trafficking dead-ends are disrupted with NZ, and these dead-ends contribute, in part, to primary ciliogenesis defects in T21 and Q21 cells. The remaining PCNT puncta likely create trafficking roadblocks that still reduce primary ciliogenesis.

### Elevated PCNT puncta and associated MTs inhibit centriolar satellite localization

To investigate the impact that enlarged PCNT puncta have on intracellular trafficking, we tested whether centriolar satellites that traffic along MTs are mislocalized with elevated HSA21 dosage. The fluorescence intensity of PCM1 was quantified in D21, T21, and Q21 cells (Fig. 4, A, B, and C). Consistent with elevated PCNT localization (Fig. 2 C), PCM1 increases around centrosomes in T21 and Q21 cells (Fig. 4 C). Though whole cell PCM1 intensity is elevated 1.3-fold in both T21 and Q21 cells, PCM1 intensity in regions surrounding centrosomes are elevated to an even greater extent (1.6- and 1.8-fold) in T21 and Q21 cells, respectively (Fig. 4, B and C). Consistent with the notion that the increased PCM1 puncta associated with PCNT puncta represent centriolar satellites, both CEP131 and CEP290 are found in the pericentrosomal crowding of T21 and Q21 cells (Fig. S4 A-C). Though CEP290 fluorescence intensities were consistently elevated in T21 and Q21 cells on average, CEP131 intensities were more variable (Fig. S4 C). Colocalization of PCNT, PCM1, CEP290, and CEP131 at cytoplasmic puncta revealed there was strong, yet variable, overlap between these molecules (Figures S4, D and E).

**Figure 4:**
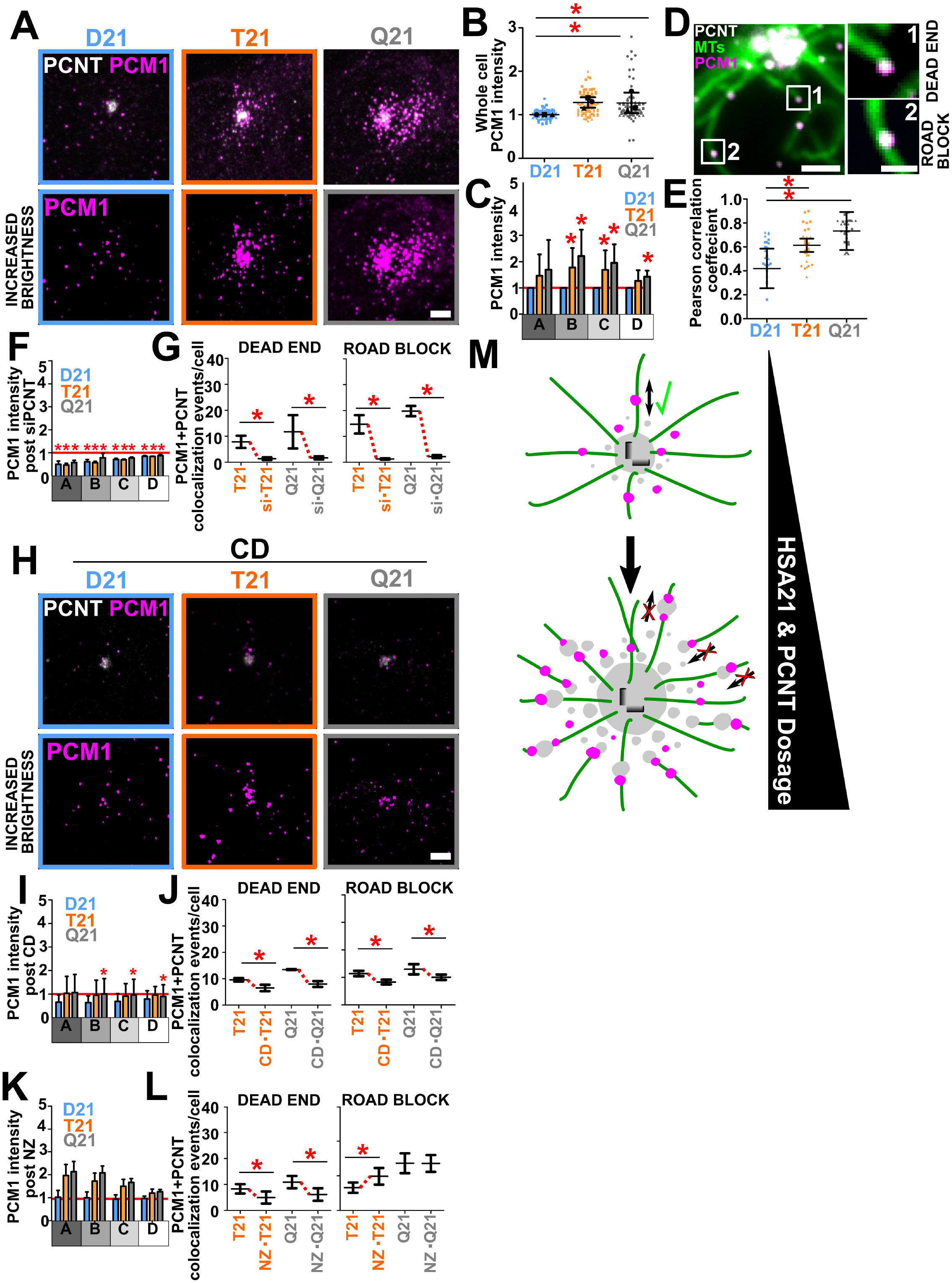
PCNT nucleated free microtubules and trafficking puncta inhibit proper trafficking of PCM1. **(A)** D21, T21 and Q21 RPE1 cells stained for PCNT (PCNT; grayscale) and PCM1 (PCM1; magenta). Brightness increased by a factor of 2.5. Scale bar, 3 μm. **(B)** D21, T21, and Q21 whole cell PCM1 fluorescence intensity. Intensity values normalized to D21 average. Statistical comparisons made to D21 averages. Mean±SD. *p < 0.05 (Table 1). **(C)** D21, T21, and Q21 cell regional binned PCM1 fluorescence intensities. Intensity values normalized to D21 average (indicated with red line). Statistical comparisons made to D21 averages. Mean±SD. *p < 0.05 (Table 1). **(D)** PCM1 colocalizes with PCNT along MTs and at MT ends. Unciliated D21 RPE1 cell stained for PCNT (PCNT; grayscale), MTs (DM1A; green) and PCM1 (PCM1; magenta). Labels are referenced in bottom panels. Scale bar, 2 μm and 0.875 μm for insets. **(E)** PCM1 and PCNT colocalization increases with Hsa21 ploidy. Mean±SD. *p < 0.05 (Table 1). **(F)** Reducing PCNT via siRNA reduces regional PCM1 intensities in D21, T21, and Q21 RPE1 cells. Red line denotes D21 siControl intensity. Statistical comparisons made to relative control intensities for each cell type. Mean±SD. *p < 0.05 (Table 1). **(G)** Reducing PCNT via siRNA reduces PCM1 colocalization with PCNT at MT ends (dead end) and along MTs (road block) in T21 and Q21 RPE1 cells. Mean±SD. *p < 0.05 (Table 1). **(H)** D21, T21 and Q21 RPE1 CD treated cells stained for PCNT (PCNT; grayscale) and PCM1 (PCM1; magenta). Brightness increased by a factor of 2.5. Scale bar, 3μm. **(I)** Cold-depolymerization reduces regional PCM1 intensities in D21, T21, and Q21 RPE1 cells. Red line denotes D21 control intensity. Statistical comparisons made to relative control intensities for each cell type. Mean±SD. *p < 0.05 (Table 1). (**J)** Cold-depolymerization reduces PCM1 colocalization with PCNT at MT ends and along MTs in T21 and Q21 RPE1 cells. Mean±SD. *p < 0.05 (Table 1). **(K)** Nocodazole does not reduce regional PCM1 intensity. Red line indicates D21 control intensity. **(L)** Nocodazole reduces PCM1 colocalization with PCNT at MT ends and increases PCM1 colocalization with PCNT along MTs in T21 cells. Mean±SD. *p < 0.05 (Table 1). **(M)** Elevated PCNT caused by Hsa21 dosage increases PCNT along MTs and increases PCNT nucleated MTs. PCNT nucleated MTs more distant from the centrosome creates trafficking dead ends, inhibiting the ability for PCM1 to traffic to the centrosome. Enlarged PCNT puncta along MTs increase trafficking roadblocks that inhibit the ability for PCM1 to traffic to and from the centrosome. The inability for proteins to efficiently traffic to and from the centrosome likely inhibits primary cilia formation. PCM1; magenta. PCNT; gray. MTs; green.

Interestingly, PCNT puncta containing both CEP131 and PCM1 or CEP131 and CEP290 were most common in D21, T21, and Q21 cells (Fig. S4 E). These results suggest that centriolar satellites are associated with the increased PCNT puncta in the pericentrosomal crowd.

To determine whether colocalization of PCNT puncta and centriolar satellites is dependent on HSA21 ploidy, we calculated Pearson correlation coefficients denoting colocalization of PCNT and PCM1 (Fig. 4, D and E). Colocalization is increased in T21 and Q21 cells compared to D21 cells. Thus, PCM1 and PCNT increasingly colocalize when PCNT is elevated. Importantly, PCM1 colocalizes with PCNT at both PCNT-nucleated free MTs (dead ends) and along MTs (roadblocks) suggesting that proper trafficking of PCM1 to and from centrosomes is disrupted with increases to either of these PCNT populations (Fig. 4 D).

To test whether increased accumulation of PCM1 peripheral to the centrosome is caused by elevated PCNT in these regions, PCM1 intensities were measured after PCNT reduction (Fig. 4 F). Indeed, reducing PCNT by siRNA also reduces PCM1 accumulation at and surrounding the centrosome. To determine whether colocalization of PCM1 and PCNT at MT ends (dead ends) or along MTs (roadblock) changed after PCNT siRNA treatment, the number of colocalization events for each population was quantified (Fig. 4 G). Importantly, colocalization of PCNT and PCM1 at both populations drastically decreases after PCNT reduction. CD, that reduces PCNT puncta and MTs, also reduces PCM1 accumulation peripheral to the centrosome in T21 and Q21 cells (Fig. 4 H and I). Moreover, colocalization of PCNT and PCM1 at both dead ends and roadblocks was reduced after CD (Fig. 4 J). These data indicate that when PCNT is elevated, PCM1 accumulates at PCNT puncta both along MTs and at free MT ends within the pericentrosomal crowding. PCNT siRNA and CD in T21 and Q21 cells both rescue the mislocalization of centriolar satellites important for primary cilia formation and function.

Consistent with NZ affecting the cell’s MT population without affecting enlarged PCNT puncta accumulation (Fig. S3 H-L), PCM1 intensity around the centrosome remained elevated in T21 and Q21 cells post NZ treatment (Fig. 4 K). Moreover, only colocalization of PCNT and PCM1 at MT ends (dead ends) was reduced after NZ treatment while PCNT and PCM1 remained colocalized along MTs (Fig. 4 L). PCNT and PCM1 colocalization along MTs (roadblock) was even increased in T21 cells and unchanged in Q21 cells after NZ treatment (Fig. 4 L). For T21 cells, this suggests a redistribution of enlarged PCNT puncta with associated PCM1 from MT ends to along MTs. For Q21 cells, this could suggest an already achieved saturation of enlarged PCNT puncta along MTs, limiting space along MTs for redistribution of puncta affected by NZ (Fig. 4 L). That there is more variability in the NZ rescue of cilia frequency than PCNT knockdown or CD suggests the remaining enlarged cytoplasmic PCNT puncta along MTs inhibit efficient ciliogenesis (Fig. S3, H, I and J). This suggests that MT dead end and trafficking roadblock populations of enlarged PCNT puncta contribute to primary cilia formation defects.

### Summary

Here, we show elevated levels of PCNT, an HSA21 resident gene upregulated in Down syndrome, increases the number of enlarged ectopic PCNT foci peripheral to the centrosome. These foci disrupt the formation of primary cilia by associating with MTs away from the centrosome, by generating MT dead-ends, and by preventing the normal distribution of the centriolar satellites required for efficient ciliogenesis by acting as roadblocks along MTs. Resetting MT distributions via depolymerization or reducing the number of enlarged PCNT foci allows for normal trafficking and for formation of primary cilia. Future work will explore which components are waylaid in the pericentrosomal crowd and what their overall dynamics are during trafficking. Moreover, we envision that dysregulated MTs and trafficking could also impact cell types like pancreatic β-cells that rely on massive changes to the interphase MT array for signaling and secretion (Zhu et al., 2015).

## Materials and Methods

### Cell Culture

Disomy 21, trisomy 21, and tetrasomy 21 hTERT-immortalized retinal pigment epithelial (RPE-1) cells were generated by Drs. Andrew Lane and David Pellman (Lane *et al*., 2014). Cells were grown in DMEM:F12 (SH30023; Cytiva) supplemented with 10% fetal bovine serum (FBS, Peak Serum; PS-FB2) and 1% Penicillin/Streptomycin at 37°C and 5% CO_2_. Cells were passaged 1:5 at ∼80-90% confluency with 0.25% Trypsin (15090-046; Gibco). Co-culture assays were performed by incubating cells in 2.5 μM CFSE (65-0850-84; eBiosciences;) in phosphate buffered saline (PBS; 1 mM KH_2_PO_4_, 155 mM NaCl, 3 mM Na_2_HPO_4_-7H_2_0, pH7.4) for eight minutes at room temperature before washing three times in complete media. Cells were then mixed and plated with unlabeled cells on 12 mm glass coverslips.

### Immunofluorescence

12 mm glass coverslips were washed in 1 M HCL heated to 50°C for 16 hours. Coverslips were then washed with water, 50%, 70%, and 95% ethanol in a sonicating water bath for 30 minutes. Coverslips were coated with collagen (C9791; Sigma) and left to air dry for 20 minutes before crosslinking under UV light for 30 minutes. Coverslips were then washed three consecutive times in PBS before addition of cells. All cells were serum starved in DMEM:F12 (SH30023; Cytiva) supplemented with 0.5% FBS (PS-FB2; Peak serum) for 24 or 48 hours prior to fixation. All cells were fixed at ∼90% confluency. Pre-permeabilized MT fixation was performed according to (Waterman-Storer and Salmon, 1997). Briefly, cells were pre-permeabilized in 0.5% TritonX-100 in PHEM (60mM Pipes, 25mM Hepes, 10 mM EGTA, 2mM MgCl2, 6.9 pH) for five minutes. Cells were then fixed with 4% paraformaldehyde / 0.5% glutaraldehyde diluted in PHEM for 20 minutes. Coverslips were quenched three times, for five minutes each, in 0.1% sodium borohydride freshly diluted in PHEM. Coverslips were then washed three times, for five minutes each, in 0.1% TritonX-100 in PHEM (PHEM-T) and stored in PHEM at 4°C until immunostaining. For non-pre-permeabilized cells, MT fixation was performed identically, but the 5-minute pre-permeabilization step was eliminated. For methanol MT fixation and all other fixations, cells were fixed in 100% methanol at -20°C for two minutes. Cells were then washed once in PBS for five minutes before a ten-minute permeabilization step in 0.5% TritonX-100 in PBS. Cells were then washed three times, for five minutes each, in PBS and stored in PBS at 4°C until immunostaining. For immunostaining, fixed cells were blocked with 0.5% bovine serum albumin (BSA, A3912-50G; Sigma) in PBS with 0.25% TritonX-100 for one hour at room temperature. Cells were incubated at 4°C overnight with primary antibodies diluted in blocking buffer. For MT immunostaining, coverslips were washed three times, for five minutes each, in PHEM-T before incubation with secondary antibodies. For all other immunostaining, coverslips were washed three times, for five minutes each, in PBS before incubation with secondary antibodies. Secondary antibodies and Hoechst33258 (1μg/mL; Invitrogen) were diluted in blocking buffer and incubated for one hour at room temperature. Coverslips were washed four times, for five minutes each, in PHEM (MT immunostaining) or PBS (all other immunostaining) before mounting to slides using Citifluor (17970-25; Electron Microscopy Sciences). Coverslips were sealed to slides using clear nail polish.

### RNAi

Human PCNT siRNA (Smart Pool) (M-012172-01-0005; Dharmacon) was transfected into cells with Lipofectamine RNAi MAX (13778100; ThermoFisher Scientific) according to the manufacturer’s protocol. Mission siRNA universal negative control #1 was used for all negative controls (SIC001-1NMOL; Sigma). All siRNAs were used at a final concentration of 25 nM. Cells were treated with siRNA in starvation media for 24 hours before fixation and subsequent immunostaining steps.

### DNA FISH

Fluorescence in situ hybridization (FISH) was performed by first washing cells in PBS and treating with 0.59% KCl for 15 minutes at 37°C. Cells were then fixed and washed three times in methanol:acetic acid 3:1 at - 20°C for five minutes each. Cells were then hybridized using a 21qter subtelomere specific probe (LPT21QG-A; Cytocell) according to the manufacturer’s protocol.

### CRISPR-Cas9

PCNT-specific crRNA (Hs.Cas9.PCNT.1.AD and Hs.Cas9.PCNT.1.AG) were designed using IDT’s online platform. Editing efficiencies of both gRNAs were confirmed by in vitro Cas9 digestion assay. For editing PCNT in T21 RPE1 cells, approximately 450,000 mycoplasma-free cells were seeded into two wells of a 6-well plate in DMEM (11995-065; Thermo Fisher Scientific) containing 10% FBS (PS-FB3; Peak Serum) and 1% Anti-Anti (100X) (15240-02; Gibco). Cells were then cultured for 24 hours at 37°C and 5% CO_2_. Cells were co-transfected with PCNT-specific RNPs that were created by complexing 1 μM Alt-R® S.p. Cas9 Nuclease V3 (1081059; IDT) with 1 μM PCNT gRNAs AD and AG, which were generated by annealing the PCNT crRNA with Alt-R^®^ CRISPR-Cas9 tracerRNA ATTO550 (1075927; IDT). Lipofectamine RNAi MAX (13778100; ThermoFisher Scientific) was used for transfection. Media was changed after 24 hours, and cells were maintained as usual (37°C and 5% CO2). The genomic DNA (gDNA) from control (non-transfected) and transfected pooled cell line was extracted and used for Polymerase Chain Reaction (PCR) to evaluate for the presence of a subpopulation with PCNT edits. Single cell clones were plated from the edited pooled populations by manually seeding approximately 1 cell into each well of 96 well plates. CloneSelect Imager (Molecular Devices; CSI8086) was used to visually select wells with single colony. These single colonies were expanded and evaluated for edit in PCNT using PCR (Fig. S1E-G).

### QIAGEN dPCR

Genetic CRISPR/Cas9 correction of PCNT copy number from 3n to 2n in trisomy 21 RPE1 cells was confirmed using the Qiagen digital PCR system (dPCR).

### Nocodazole and Cold-Depolymerization

Cold-depolymerization (CD) experiments were performed by first placing cells in starvation media for 24 hours. At 24 hours, cells were placed on ice for 10 or 20 minutes at 4°C. For CD MT stability assays, cells were fixed directly following a 10-minute CD. For 20-minute CD, cells were returned to incubation at 37°C for additional 24 hours prior to fixation. All control cells were left at 37°C. Nocodazole treatments were performed by first placing cells in starvation media for 24 hours. At 24 hours, cells were treated with 1 μM nocodazole for 60 minutes at 37°C. After 60 minutes, coverslips were washed five times, for 5 minutes each, in starvation media. After washes, cells were returned to incubation at 37°C for additional 24 hours prior to fixation. Control cells were treated with the same volume of Dimethyl Sulfoxide (DMSO, D2650-100ML; Sigma) for 60 minutes at 37°C.

### Generation of tetracycline-inducible mNeon-EB3 RPE-1 D21, T21, and Q21 cell lines

Tetracycline-inducible lentiviral stable RPE-1 cell lines were generated according to methods outlined in (Sankaran et al., 2020). Briefly, HEK293T cells were transfected with the tetracycline-inducible mNeon-EB3 construct and lentivirus packaging plasmids using Lipofectamine 2000 (11668-027; Invitrogen). HEK293T media containing virus was collected and added to target cells in the presence of 2 μg/mL polybrene. Target cells were incubated with viral media for 24 hours. After 24 hours, new viral media was added to target cells and target cells were left to incubate additional 24 hours. Transduced cells were selected using 10 μg/mL puromycin for three days. Cells were flow-sorted to isolate EB3-mNeon positive clones and induced overnight with 0.125 μg/mL doxycycline before fixation.

### Fluorescence microscopy

Widefield images were acquired with a Nikon Eclipse Ti-E microscope (Nikon) equipped with a 100x Plan Apochromat objective (NA 1.40) and an Andor Xyla 4.2 scientific CMOS camera (Andor). Nikon NIS Elements imaging software was used for widefield image acquisition. Confocal images were acquired with a Nikon Eclipse Ti inverted microscope stand equipped with a 100x Plan Apochromat objective (NA 1.45), Andor iXon X3 camera, and CSU-X1 (Yokogawa) spinning disk. SIM images were acquired using a Nikon SIM (N-SIM) with a Nikon Ti2 (Nikon Instruments; LU-N3-SIM) microscope equipped with a 100× SR Apo TIRF, NA 1.49 objective. Images were captured using a Hamamatsu ORCA-Flash 4.0 Digital CMOS camera (C13440) with 0.1-μm Z step sizes. Raw SIM images were reconstructed using the image slice reconstruction algorithm (NIS Elements). Slidebook 6 digital microscopy software was used for confocal image acquisition. Image acquisition times were kept constant within a given experiment and ranged from 30 to 500 msec. All images were acquired at room temperature. Identical acquisition settings were used for quantitatively compared images. Confocal imaging was used to assess colocalization of two proteins. All other quantification was done using widefield acquired images. Images in Figures 3C, 4A, 4D, 4H, S4A, S4B, and S4D were acquired using confocal microscopy. Images in Figures 3H are SIM images. All other figure images were acquired using widefield microscopy. All images presented in figures are max projections, except for images showing MT colocalization with PCNT/PCM1. These images were selected from a single z-plane.

### Fluorescence quantification

To quantify the presence of cilia, cilia structures were labeled with ARL13B, 6-11-B1 (acetylated tubulin) or DM1A (*α*-tubulin) primary antibodies. Radial fluorescence intensity analysis was completed using the Radial Profile Extended ImageJ plugin. Briefly, this analysis plots average fluorescence intensity as a function of distance from the user identified centroid. Whole cell microtubule density analysis was performed by outlining cell boundaries (defined by MTs) in ImageJ. Free MTs are defined as being traceable and having two clear ends (identified by moving through slices of the z-stack). To evaluate colocalization of two proteins, ROIs with radius 10 μm were centered over the centrosome. Pearson correlation coefficients were recorded for two proteins of interest within the ROI using ImageJ plugin Coloc2. To quantify PCNT puncta ratios, ImageJ particle analysis was performed on 10 μm radius ROIs centered over the centrosome of cells. All other fluorescence intensity analyses were performed by recording the integrated density of defined ROIs in ImageJ. All intensity analysis was performed on max projected images.

### Statistics

Data were analyzed and organized with Microsoft Excel and GraphPad Prism. Data center values represent averages while error bars represent standard deviation. All experiments utilized at least three biological replicates. The D’Agostino and Pearson omnibus normality test was used to test for normality of data. A Student’s two-tailed unpaired t-test was used to test significance between two normal unpaired distributions. The Mann-Whitney U-test was used to test significance between two non-normal unpaired distributions. Differences between distributions were considered statistically significant if p-values were less than 0.05. P-values, sample sizes, and statistical tests used for samples are denoted in Table 1. % CV was calculated by dividing the standard deviation of a distribution by the mean of the distribution and multiplying by 100.

## Abbreviations

PCNT: Pericentrin
MT: microtubule
HSA21: Human somatic autosome 21
DS: Down syndrome

## Acknowledgements

We thank Drs. Andrew Lane and David Pellman for generously providing the HSA21 RPE1 cell lines; A. Soh for ImageJ code contributions; and Drs. J. Black and L. Porritt for assistance with FISH experiments. The Pearson lab for discussions. This research was funded by R01 GM 138415 and R35 GM 140813 to CGP, the Linda Crnic Institute for Down Syndrome, and the Global Down Syndrome Foundation. The authors declare no competing financial interests.

## Supplemental Figure Legends

**Figure S1:**
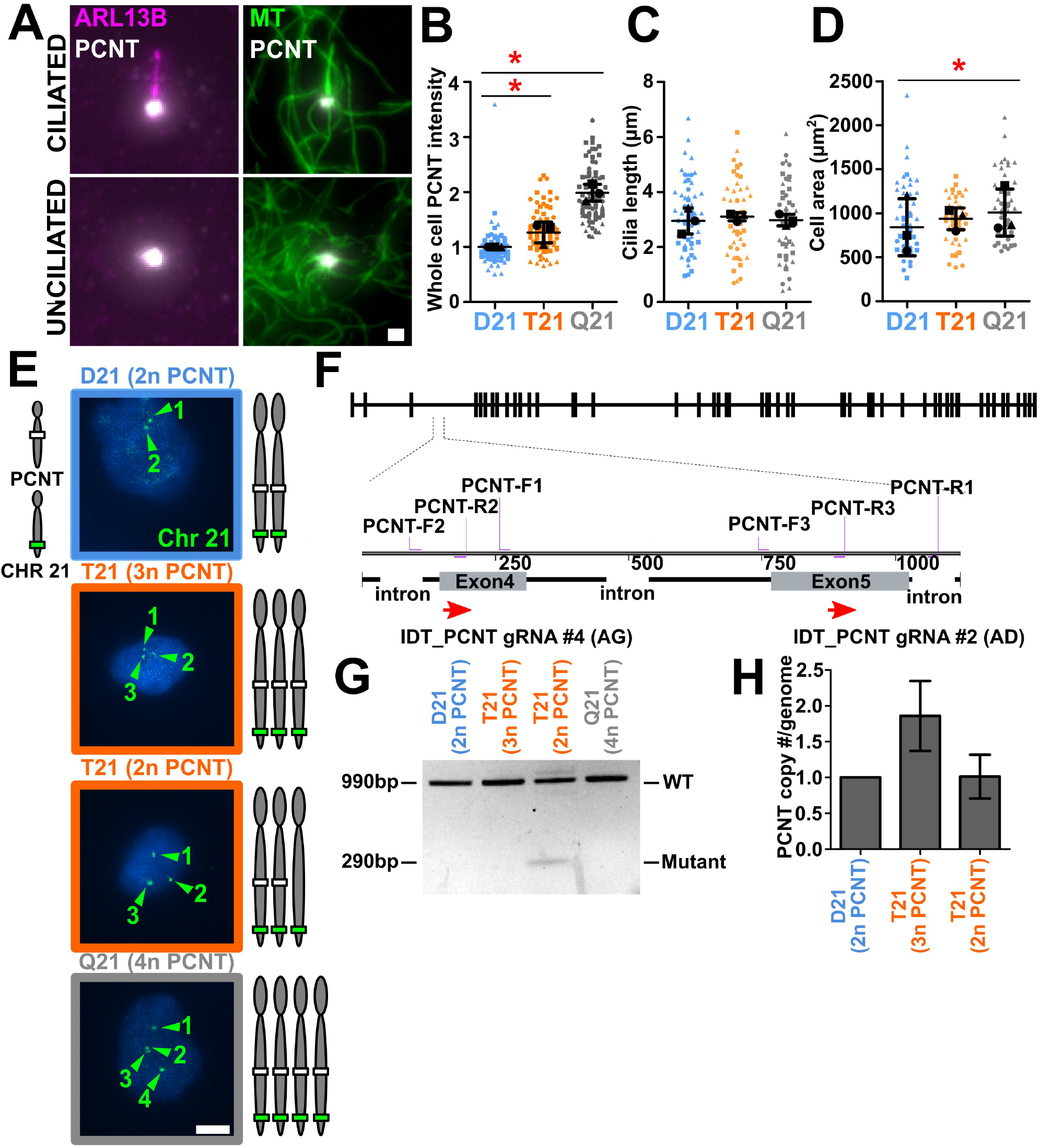
**(A)** Left, representative ciliated and unciliated cell stained for centrosomes (PCNT; grayscale) and cilia (ARL13B; magenta). Right, representative ciliated and unciliated cell stained for centrosomes (PCNT; grayscale) and cilia (DM1A; MT; green). Scale bar, 1 μm. **(B)** T21 and Q21 RPE1 cells have more PCNT. Intensities normalized to D21 averages. Mean±SD. *p < 0.05 (Table 1). **(C)** Cilia length is unchanged between D21, T21, and Q21 RPE1 cells. Mean±SD. **(D)** Cell area increases in Q21 RPE1 cells. Areas are normalized to D21 averages. Mean±SD. *p < 0.05 (Table 1). **(E)** CRISPR-Cas9 knockout of one allele of PCNT in T21 RPE1 cells (T21 (2n PCNT)) does not alter chromosome 21 copy number (HSA21 subtelomere probe; green). Scale bar, 5 μm. **(F)** Cas9 cleavage sites are located in PCNT Exon 4 and Exon 5. PCR using PCNT-F2/R1 oligos was used to verify deletion of the gene segment spanning the two Cas9 cleavage sites. **(G)** PCR verification of genetic ablation of a single PCNT allele in T21 cells using PCNT-F2/R1 oligos. WT PCNT expected band size: 990 bp. Edited PCNT expected band size: 290 bp. **(H)** QIAGEN dPCR verification of PCNT copy number / genome.

**Figure S2:**
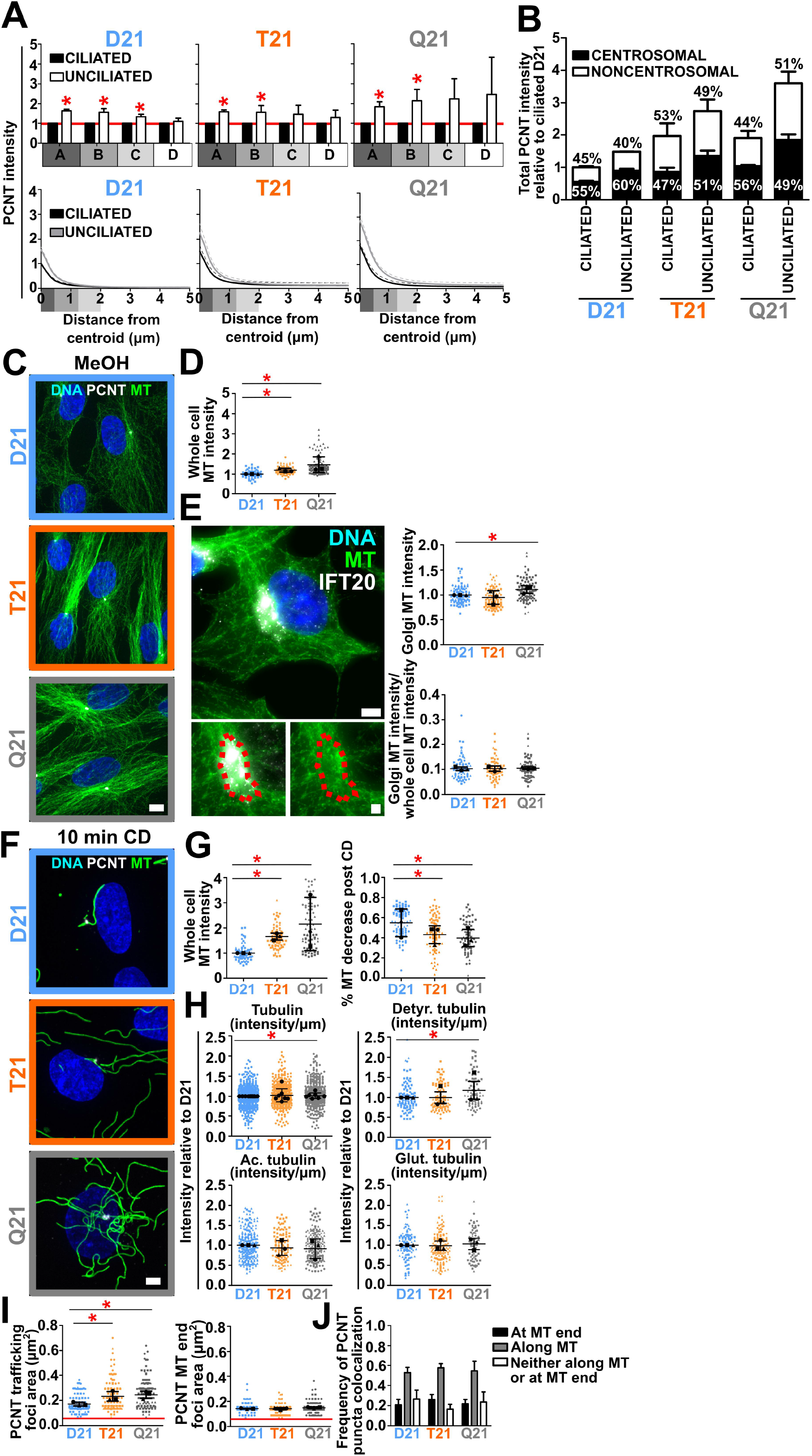
**(A)** Top, D21, T21, and Q21 cell regional binned PCNT fluorescence intensities. Intensity values normalized to ciliated cell average within each ploidy level (indicated with red line). Statistical comparisons made to ciliated averages. Bottom, D21, T21, and Q21 radial PCNT fluorescence intensities in ciliated and unciliated cells. Intensity normalized to ciliated D21 centrosome intensity. Mean±SD. *p < 0.05 (Table 1). **(B)** PCNT fluorescence in a 5 μm radius in ciliated and unciliated D21, T21, and Q21 cells. Relative fractional intensity contributions of centrosomal and non centrosomal PCNT to the total PCNT intensity is indicated. Intensity was defined as centrosomal if the fluorescence intensity fell within 0 - 1.2 µm of the centriole of the centrosome. Intensity was defined as noncentrosomal if fluorescence intensity fell within 1.2 - 5 µm of the centrosome. **(C)** D21, T21 and Q21 RPE1 cells stained for MTs (DM1A; green), PCNT (PCNT; grayscale), and DNA (Hoescht33258; blue). MTs in these images were fixed with methanol. Scale bar, 4 μm. **(D)** D21, T21, and Q21 whole cell methanol fixed MT fluorescence intensities. Intensity values normalized to D21 average. Statistical comparisons made to D21 averages. Mean±SD. *p < 0.05 (Table 1). **(E)** RPE1 cell labeled for golgi (IFT20; grayscale), MTs (DM1A; green), and DNA (Hoescht33258; blue). For golgi MT quantification, golgi boundaries were outlined (shown) and MT intensities within outline were calculated. Top, golgi MT intensity is increased in Q21 RPE1 cells. Bottom, golgi MT contributions to whole cell MT intensities are unchanged in D21, T21, and Q21 cells. Intensity values normalized to D21 average. Statistical comparisons made to D21 averages. Mean±SD. *p < 0.05 (Table 1). **(F)** 10-minute CD D21, T21 and Q21 RPE1 cells stained for MTs (DM1A; green), PCNT (PCNT; grayscale), and DNA (Hoescht33258; blue). **(G)** Left, D21, T21, and Q21 10-minute CD MT fluorescence intensities. Right, whole cell MT intensity reduction post 10-minute CD is reduced in T21 and Q21 RPE1 cells. Statistical comparisons made to D21 averages. Mean±SD. *p < 0.05 (Table 1). **(H)** D21, T21, Q21 intensity/μm values for tubulin post-translational modifications. Mean±SD. *p < 0.05 (Table 1). Tubulin, acetylated tubulin, glutamylated tubulin, and detyrosinated tubulin intensities were quantified. Intensity values normalized to D21 average. Statistical comparisons made to D21 averages. Mean±SD. *p < 0.05 (Table 1). **(I)** Left, area of PCNT foci along MTs are larger than 0.05 μm^2^ (indicated with red line) and increase with Hsa21 ploidy. Bottom, area of PCNT foci at MT minus ends are larger than 0.05 μm^2^ (indicated with red line). Mean±SD. *p < 0.05 (Table 1). Along MT, n=113 D21, 148 T21, 147 Q21. MT end, n=151 D21, 150 T21, 150 Q21. **(J)** 78% of PCNT puncta, on average, localize to pre-permeabilized MTs in D21, T21, and Q21 RPE1 cells.

**Figure S3:**
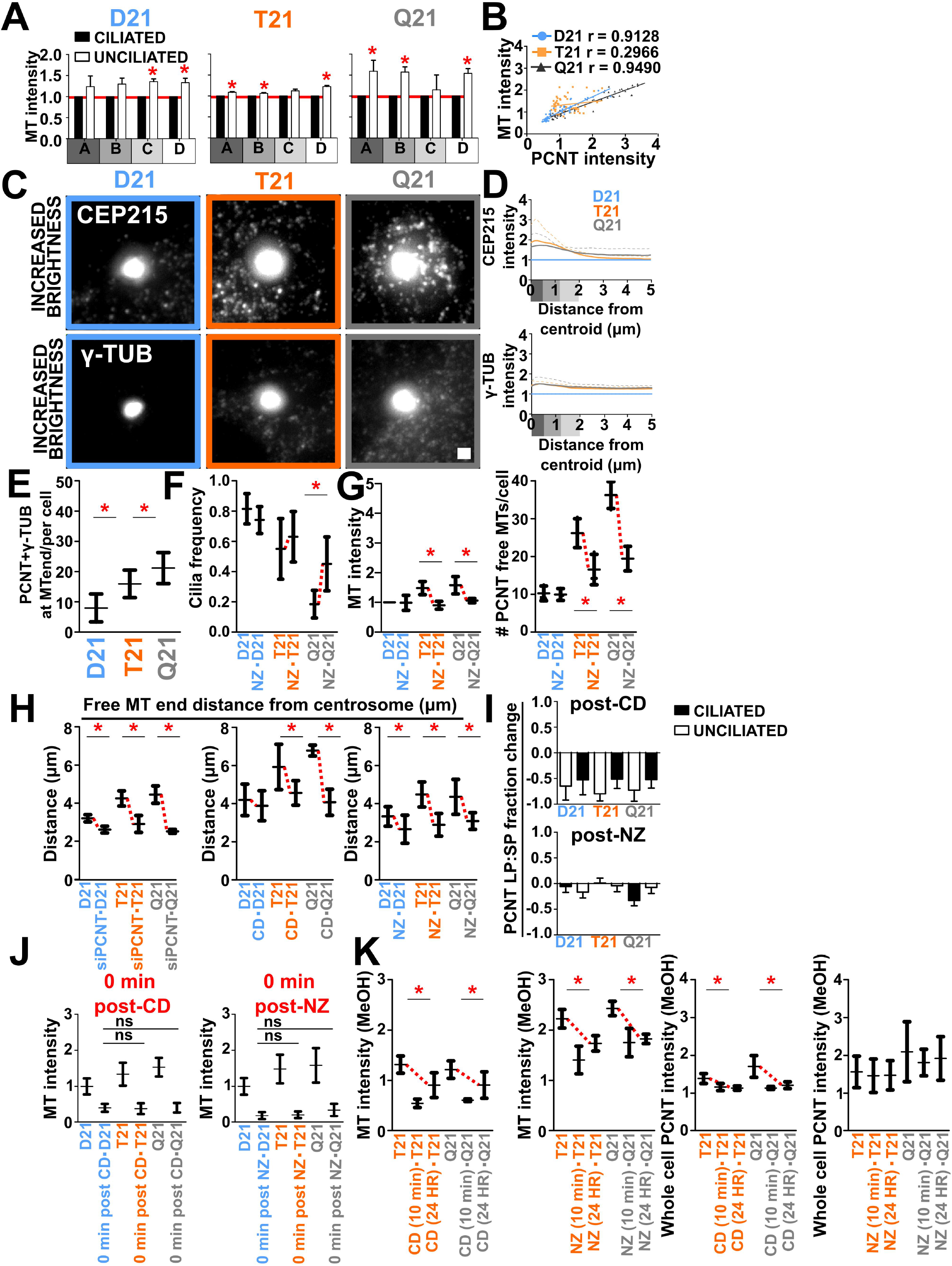
**(A)** D21, T21, and Q21 cell regional binned MT fluorescence intensities. Intensity values normalized to ciliated cell average within each ploidy level (indicated with red line). Statistical comparisons made to ciliated averages. Mean±SD. *p < 0.05. **(B)** MT and PCNT intensity correlation in D21, T21, and Q21 RPE1 cells. D21 r = 0.9128, T21 r = 0.2966, Q21 r = 0.9490 (Table 1). **(C)** PCNT interacting partners, CEP215 and *γ*-tubulin, increase with Hsa21 ploidy. Top panels, D21, T21, and Q21 cells stained for CEP215 (CEP215; grayscale). Middle panels, D21, T21, and Q21 cells stained for *γ*-tubulin (GTU88; grayscale). Scale bar 1μm. **(D)** Top, T21, and Q21 radial CEP215 fluorescence intensities normalized to D21 intensity (indicated with blue line). Bottom, T21, and Q21 radial *γ*-tubulin fluorescence intensities normalized to D21 intensity (indicated with blue line). **(E)** PCNT and *γ*-tubulin colocalization at MT ends increases with HSA21 ploidy. Mean±SD. *p < 0.05 (Table 1). **(F)** Nocodazole (NZ) treatment increases cilia frequencies in Q21 RPE1 cells. Mean±SD. *p < 0.05 (Table 1). **(G)** Left, NZ treatment reduces MT intensities in T21 and Q21 RPE1 cells. Intensity values normalized to D21 average. Data points represent 5μm binned radial MT fluorescence intensities. Right, NZ treatment reduces PCNT nucleated MTs in T21 and Q21 RPE1 cells. Mean±SD. *p < 0.05 (Table 1). **(H)** Left, reducing PCNT via siRNA reduces distance of PCNT nucleated free MTs from centrosome in D21, T21, and Q21 RPE1 cells. Middle, cold-depolymerization reduces distance of PCNT nucleated free MTs from centrosome in D21, T21, and Q21 RPE1 cells. Right, NZ treatment reduces distance of PCNT nucleated free MTs from centrosome in D21, T21, and Q21 RPE1 cells. Mean±SD. *p < 0.05 (Table 1). **(I)** Cold-depolymerization reduces mean PCNT LP:SP ratios in T21 and Q21 RPE1 cells while NZ treatment does not. Mean±SD. **(J)** Left, 60-80% of whole cell MTs in D21, T21, and Q21 RPE1 cells are depolymerized 0 minutes post 20-minute CD. Right, 75-85% of whole cell MTs in D21, T21, and Q21 RPE1 cells are depolymerized 0 minutes post NZ treatmeant. **(K)** Left two graphs, methanol fixed whole cell MT intensities appear to reach their growth capacity 10 minutes after CD or NZ treatment. Right two graphs, whole cell PCNT is reduced 10 minutes following CD and remains reduced 24 hours post CD. Whole cell PCNT is unchanged post NZ treatment. Intensity values normalized to D21 averages. Statistical comparisons made to D21 averages. Mean±SD. *p < 0.05 (Table 1).

**Figure S4:**
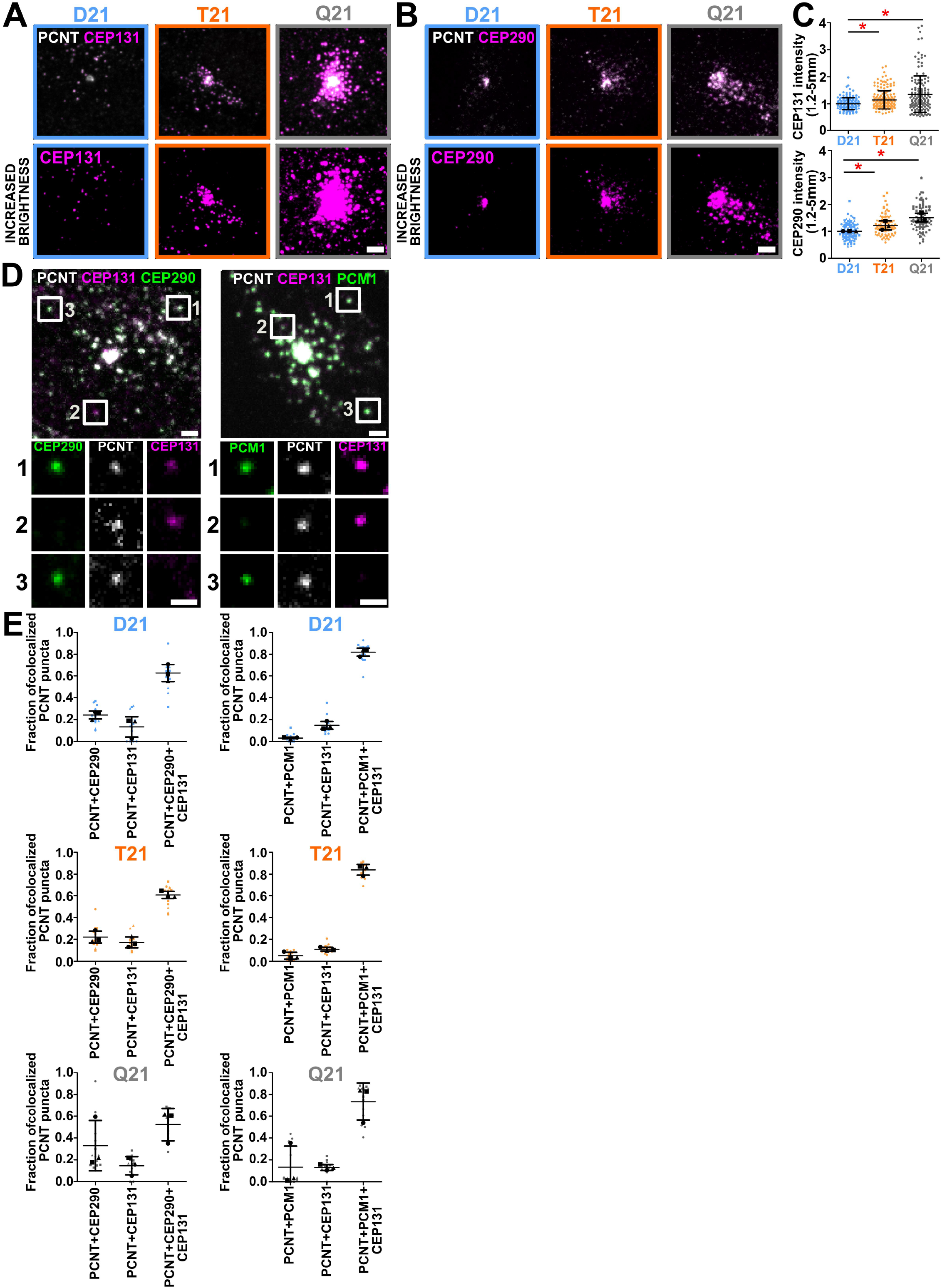
**(A)** D21, T21 and Q21 RPE1 cells stained for PCNT (PCNT; grayscale) and CEP131 (CEP131; magenta). Brightness increased by a factor of 3. Scale bar, 3 μm. **(B)** D21, T21 and Q21 RPE1 cells stained for PCNT (PCNT; grayscale) and CEP290 (CEP290; magenta). Brightness increased by a factor of 3. Scale bar, 3 μm. **(C)** Top, binned CEP131 fluorescence intensity (1.2-5μm away from the centrosome) increases in T21 and Q21 RPE1 cells. Bottom, binned CEP290 fluorescence intensity (1.2-5μm away from the centrosome) increases in T21 and Q21 RPE1 cells. Intensity values normalized to D21 averages. Statistical comparisons made to D21 averages. Mean±SD. *p < 0.05 (Table 1). **(D)** Left, RPE1 cell stained for PCNT (PCNT;grayscale), CEP131 (CEP131;magenta), and CEP290 (CEP290; green). Right, RPE1 cell stained for PCNT (PCNT;grayscale), CEP131 (CEP131;magenta), and PCM1 (CEP290; green). Insets show examples of colocalization quantified in (E). Scale bars, 1.5 μm and 1μm for insets. **(E)** Colocalization of PCNT puncta with CEP290, PCM1, and CEP131 in D21, T21, and Q21 RPE1 cells.

